# SARS-CoV-2 ORF3c impairs mitochondrial respiratory metabolism, oxidative stress and autophagic flow

**DOI:** 10.1101/2022.10.04.510754

**Authors:** Alessandra Mozzi, Monica Oldani, Matilde E. Forcella, Chiara Vantaggiato, Gioia Cappelletti, Chiara Pontremoli, Francesca Valenti, Diego Forni, Mara Biasin, Manuela Sironi, Paola Fusi, Rachele Cagliani

## Abstract

Coronaviruses encode a variable number of accessory proteins that play a role in host-virus interactions, in the suppression of immune responses, or in immune evasion. Accessory proteins in SARS-CoV-2 consist of at least twelve viral proteins whose roles during infection have been extensively studied. Nevertheless, the role of the ORF3c accessory protein, an alternative open reading frame of ORF3a, has remained elusive. Herein, we characterized ORF3c in terms of cellular localization, host’s antiviral response modulation, and effects on mitochondrial metabolism. We show that ORF3c has a mitochondrial localization and alters mitochondrial metabolism, resulting in increased ROS production, block of the autophagic flux, and accumulation of autophagosomes/autolysosomes. Notably, we also found that ORF3c induces a shift from glucose to fatty acids oxidation and enhanced oxidative phosphorylation. This is similar to the condition observed in the chronic degenerative phase of COVID-19. Altogether these data suggest that ORF3c could be a key protein for SARS-CoV-2 pathogenesis and that it may play a role in disease progression.

## INTRODUCTION

The ongoing COVID-19 pandemic, which is caused by a newly emerged coronavirus (SARS-CoV-2), has to date resulted in more than 6.4 million deaths worldwide (https://covid19.who.int/). Although vaccines have been demonstrated to be highly efficient in preventing severe disease presentation and mortality (Evans & Jewell, 2021), the emergence of new variants suggests the need for a deeper understanding of SARS-CoV-2 pathogenetic mechanisms, to improve prevention and treatment (Chakraborty et al, 2020). COVID-19 involves two stages, with different metabolic features (Shenoy, 2020). A first hyper-inflammatory phase, characterized by increased aerobic glycolysis (Warburg effect), increased oxygen consumption, elevated ATP production, and hyperglycemia, occurs as the host tissues react to the virus by increasing energy production and by activating the innate immune response. This is the phase which often culminates with the cytokine storm (Fitzpatrick, 2019; Singer, 2014). A second hypo-inflammatory, immune-tolerant phase is characterized by decreased oxygen consumption, resumption of mitochondrial respiration and ATP production, as well as by increased fatty acid oxidation (Vachharajani & McCall, 2020; Wang et al, 2018).

SARS-CoV-2 is an enveloped virus consisting of a positive-sense, single-stranded RNA genome of about 30 kb (Gordon et al, 2020b; Wu et al, 2020). Two overlapping ORFs, ORF1a and ORF1b, are translated from the positive-strand genomic RNA and generate continuous polypeptides, which are cleaved into a total of 16 nonstructural proteins (NSPs). The remaining genomic regions encode four structural proteins - spike (S), envelope (E), membrane (M) and nucleocapsid (N) - and six annotated accessory proteins (ORF3a, 6, 7a, 7b, 8 and 10; reference GenBank ID: NC_045512.2). Moreover, studies that aimed to evaluate the coding capacity of SARS-CoV-2 identified several unannotated viral accessory ORFs, including several alternative open reading frames within ORFs S (ORF2d), N (ORF9b, ORF9c), and ORF3a (ORF3b, ORF3c, ORF3d) (Finkel et al, 2021).

Previously, protein-protein interaction data between SARS-CoV-2 proteins and cellular molecules were obtained using different methods, such as affinity purification, proximity labeling-based strategy, and yeast two-hybrid system (Chen et al, 2021b; Davies et al, 2020; Gordon et al, 2020a; Gordon et al, 2020b; Stukalov et al, 2021; Wu et al, 2020). These host-virus interactome analyses uncovered several human proteins that physically associate with SARS-CoV-2 proteins and that may participate in the virus life cycle, infection, replication and budding. Among these, interactions with mitochondrial proteins seem to be particularly abundant (Gordon et al, 2020a; Gordon et al, 2020b; Stukalov et al, 2021). In line with these findings, recent studies suggested the involvement of mitochondria in SARS-CoV-2 infection as a hallmark of disease pathology (Edeas et al, 2020; Guzzi et al, 2020; Kloc et al, 2020; Singh et al, 2020). Moreover, recent evidence revealed alterations of mitochondrial dynamics in COVID-19 patients, associated with increased fusion, and inhibition of mitochondrial fission (Holder & Reddy, 2021).

In this respect, the study of accessory proteins with mitochondrial localization is of great importance to identify therapeutic targets and to understand the mechanisms of SARS-CoV-2-induced disease (Yan et al, 2022). Indeed, although accessory proteins are considered non-essential for coronavirus replication, accumulating evidence demonstrates that they are critical to virus-host interaction, affecting host innate immunity, autophagy, and apoptosis, as well as contributing significantly to pathogenesis and virulence (Redondo et al, 2021). For instance, the ORF9b protein, which localizes to mitochondria, antagonizes type I and III interferons by targeting multiple innate antiviral signaling pathways (Han et al, 2021). Another mitochondrial accessory protein, ORF10, inhibits the cell innate immune response inducing mitophagy-mediated MAVS degradation (Li et al, 2022a).

A notable exception among SARS-CoV-2 accessory proteins is accounted for by ORF3c, which has remained uncharacterized and under-investigated. The ORF3c protein has been predicted to be encoded by sarbecoviruses (a subgenus of betacoronaviruses) only (Firth, 2020; Jungreis et al, 2021), including SARS-CoV-2, SARS-CoV, and bat coronavirus RaTG13 (one of the bat betacoronavirus most closely related to SARS-CoV-2 (Zhou et al, 2020)). Analysis of the conservation of ORF3c in sarbecoviruses, together with ribosome-profiling data, strongly suggest that ORF3c is a functional protein (Cagliani et al, 2020; Finkel et al, 2021; Firth, 2020; Jungreis et al, 2021).

Herein, we report the first investigation of the effect of ORF3c on cellular innate immune responses, autophagy, and lung cell mitochondrial metabolism.

## RESULTS

### ORF3c protein structure

SARS-CoV-2 ORF3c (also known as ORF3h) is a 41 amino acid (aa) protein encoded by an alternative open reading frame within the ORF3a gene (Cagliani et al, 2020; Firth, 2020; Jungreis et al, 2021). It is highly conserved in sarbecoviruses with 90% identity with SARS-CoV and 95% with bat-CoV RaTG13 (Fig. 1A). This latter was isolated from horseshoe bats (*Rhinolophus affinis*), a putative reservoir host, and it is one of the bat viruses most closely related to SARS-CoV-2 (Zhou et al, 2020).

**Figure 1.**
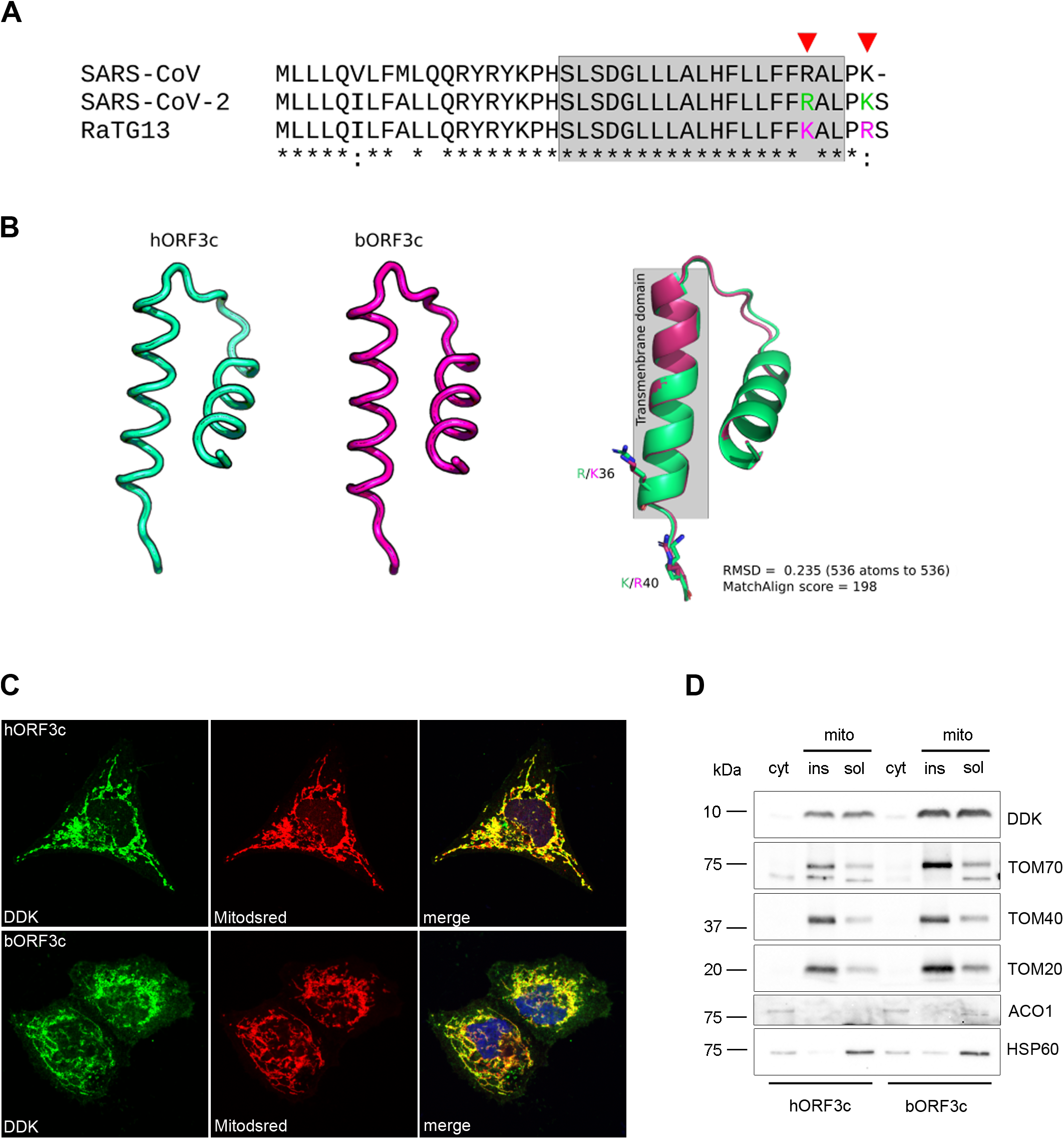
ORF3c localizes to the mitochondria. **(A)** ClustalW alignment of SARS-CoV-2 ORF3c (hORF3c), bat CoV RaTG13 ORF3c (bORF3c) and SARS-CoV ORF3c proteins. Transmembrane domains predicted with Phobius (https://phobius.sbc.su.se/) are in gray. The amino acid positions 36 and 40 specific for hORF3c and bORF3c are marked in green and magenta, respectively. **(B)** Protein structures of hORF3c and bORF3c modeled with the RoseTTAFold software. Superimposition of the two structures is also reported and visualized with PyMOL. **(C)** Mitochondrial localization of ORF3c proteins. HeLa cells were cotransfected with pDsRed2-Mito vector and pCMV6 hORF3c or bORF3c. Twenty-four hours later, cells were fixed and immunostained with antibodies against the DDK tag. Co-localization (yellow) of ORF3c (green) with mitochondria (red) is shown in the merged images. Nuclei were counterstained with DAPI (blue). Scale bar: 10 μm. **(D)** HeLa cells transiently expressing hORF3c or bORF3c were lysed and total cell extracts were subjected to cellular fractionation. Aliquots of cytosolic and mitochondrial (soluble/insoluble) fractions were analysed by SDS–PAGE and Western blotting. hORF3c and bORF3c were detected using an anti-DDK antibody. Antibodies directed against the cytosolic protein aconitase 1 (ACO1), the outer mitochondrial membrane translocase subunits TOM20, TOM40 and TOM70, and the mitochondrial matrix heat shock protein 60 (HSP60) were used as markers of the specific cellular compartment/organel.

As previously reported, ORF3c has a predicted highly conserved transmembrane domain (Cagliani et al, 2020) (Fig. 1A), which suggests interactions within the lipid bilayer (Redondo et al, 2021). However, other protein domains have not been described and the protein structure is not available. We thus modeled the structure of the SARS-CoV-2 and bat CoV RaTG13 ORF3c proteins with the RoseTTAFold software using the deep-learning algorithm (Baek et al, 2021). ORF3c structure prediction revealed a tridimensional architecture composed of two short alpha-helices (α1 and α2) connected by a loop region (Fig. 1B). The α2 helix corresponds to the predicted transmembrane region. SARS-CoV-2 and RatG13 ORF3c proteins differ only in two amino acids: R36K (in the predicted transmembrane domain) and K40R (Fig. 1A). Structural superposition revealed good conservation of the global protein architecture between the two models (Fig. 1B), suggesting that amino acid differences between human and bat ORF3c proteins do not result in conformational changes.

### SARS-CoV-2 ORF3c localizes to the mitochondria

ORF3c subcellular localization was investigated by confocal microscopy. In particular, 123 bp sequences corresponding to the ORF3c of SARS-CoV-2 and RaTG13 (hereafter hORF3c and bORF3c, respectively) were cloned into a mammalian expression vector (pCMV6) in frame with the DDK (FLAG) tag. HeLa cells were transiently transfected with the vectors expressing hORF3c and bORF3c and stained with anti-DDK antibody to detect the viral protein, as well as with antibodies against specific markers of the endoplasmic reticulum, Golgi, lysosomes or early endosomes. For the staining of mitochondria, cells were transfected with the pDsRed2-Mito vector. Immunofluorescence analysis revealed that both hORF3c and bORF3c strongly co-localized with mitochondria (Fig. 1C), as already reported for other SARS-CoV-2 accessory proteins, such as ORF9b (Jiang et al, 2020). This latter was previously shown to directly interact with the outer mitochondrial membrane protein TOM70 (translocase of outer membrane 70) (Jiang et al, 2020), which forms the translocon complex with other TOM proteins (Eaglesfield & Tokatlidis, 2021). We found that hORF3c and bORF3c proteins colocalize with TOM70 and TOM20 (Suppl Fig. S1A). However, a direct interaction between the two ORF3c proteins and TOM complex (TOM70, TOM20 and TOM40) was excluded, as indicated by immunoprecipitation analysis (Suppl Fig.S1B). The mitochondrial localization of both ORF3c proteins was also confirmed in the two lung cell lines A549 and HSAEC1 (Suppl Fig S1C), deriving from lung carcinomatous tissue and normal lung tissue, respectively.

Fractionation analysis in HeLa cells confirmed that hORF3c and bORF3c were almost exclusively found in the mitochondria, in both soluble and insoluble (membrane) fractions (Fig. 1D). Altogether, these data indicate that ORF3c localizes in the mitochondria and suggest that, at least partially, the protein product of ORF3c localizes on mitochondrial membranes. Further studies are required to clarify whether mitochondrial membrane binding of ORF3c occurs via the predicted transmembrane domain and/or via interaction with host mitochondrial proteins.

### The SARS-CoV-2 ORF3c protein induces an increase in mitochondrial respiratory metabolism, a reduction in glycolysis and a metabolic shift towards dependency on fatty acids

Because the ORF3c protein localizes in the mitochondria, we investigated whether it acts by modifying mitochondrial metabolism. Mitochondrial functionality of HSAEC1 cells transfected with hORF3c and bORF3c were investigated through Agilent Seahorse XF Mito Stress analysis. In particular, results obtained by measuring real-time oxygen consumption rate (OCR) showed that hORF3c protein increases both basal and maximal respiration, as well as mitochondrial ATP synthesis (Figure 2A and C). However, this was not matched by an increase in glycolysis, since no differences were observed among extra-cellular acidification rate (ECAR) profiles (Fig. 2B). An increase in both maximal respiration and spare respiratory capacity was observed in HSAEC1 cells overexpressing the RaTG13 ORF3c protein, while the basal respiration increase was not found statistically significant (Fig. 2C). Moreover, cells transfected with hORF3c or bORF3c showed a slight increase in oxygen consumption after oligomycin addition (Fig. 2C). Although this result may be correlated to mitochondrial uncoupling, the mitochondria of cells overexpressing viral ORF3c proteins are not uncoupled (Suppl Fig. S2A). Mitochondrial Δψ, measured using a DiOC6 (3,3’-dihexyloxacarbocyanine iodide) fluorescent probe, was found to be more negative in both transfected cells compared to the control (Fig. 2D), suggesting oxidative phosphorylation hyperactivation.

**Figure 2.**
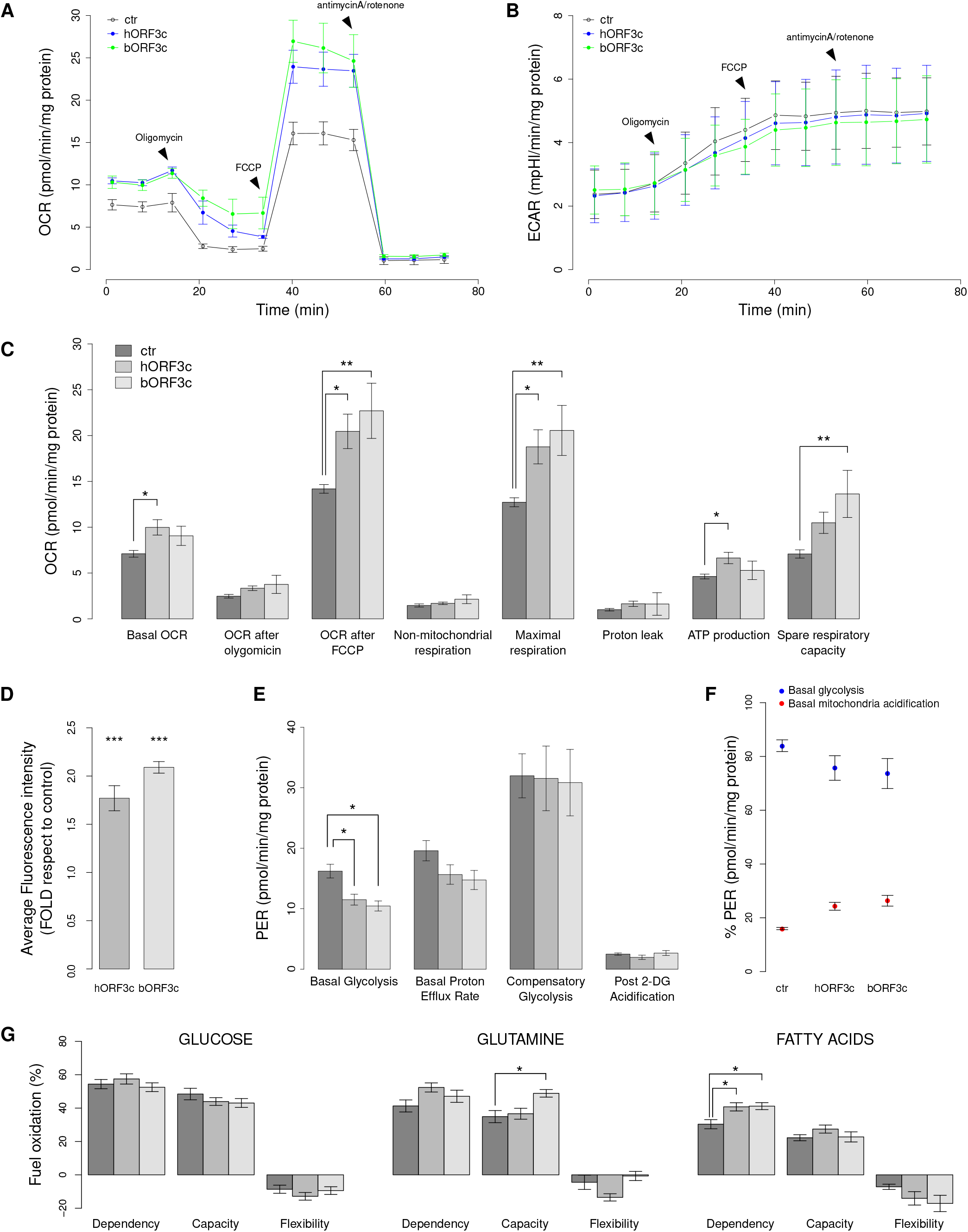
ORF3c modifies mitochondrial metabolism. **(A)** Seahorse mitostress analysis in HSAEC1 cells transfected with hORF3c or bORF3c. OCR traces are expressed as pmoles O2/min/mg proteins. The arrows indicate the time of oligomycin, FCCP and antimycinA/rotenone addiction. The OCR profile is representative of three independent experiments. **(B)** ECAR traces are expressed as mpH/min/mg proteins. The arrows indicate the time of oligomycin, FCCP and antimycinA/Rotenone addition. The ECAR profile is representative of three independent experiments. **(C)** Bars (mean ± SEM obtained in three independent experiments) indicate the values at points 3 (basal OCR), 6 (OCR after oligomycin), 9 (OCR after FCCP) and different parameters related with mitochondrial function (non-mitochondrial respiration, maximal respiration, proton leak, ATP production, spare respiratory capacity). **(D)** Analysis of mitochondrial Δψ. After transfection, cells were incubated with 40 nM DiOC6 and the level of fluorescence was evaluated. Results are representative of three independent experiments. **(E)** Seahorse glycolytic analysis. Analysis of different parameters related with glycolysis (basal glycolysis, basal proton efflux rate, compensatory glycolysis, post-2DG acidification). **(F)** Proton Efflux Rate (PER) due to glycolysis and to oxidative phosphorylation. **(G)** Evaluation of mitochondrial fuel oxidation in HSAEC1 cells transfected with ORF3c from either human or bat SARS-CoV-2. Glucose, glutamine and long-chain fatty acids mitochondrial fuel oxidation dependency, capacity and flexibility. Bars indicate the mean ± SEM obtained in three independent experiments. Statistical significance: * p < 0.05, ** p < 0.01, *** p < 0.001 (Dunnett’s test).

In the XF Seahorse Glycolysis Rate Assay we observed a decrease in the level of basal glycolysis in transfected cells, as well as a decreasing trend in the basal proton efflux rate (PER) (Fig. 2E). PER percentage allows to distinguish between basal mitochondria acidification, due to CO_2_ release, and glycolytic acidification, due to lactic acid production. The overexpression of each ORF led to an increase of the PER derived from mitochondria and a decrease in glycolytic PER (Fig. 2F). In accordance, the activity of lactate dehydrogenase (LDH) did not significantly increase after transfection (Supplementary Fig. S2B), suggesting that pyruvate is predominantly used in the Krebs cycle.

Subsequently, mitochondria dependence on various substrates was investigated through the Seahorse Mito Fuel Flex Test Kit. In particular, the dependency, the capacity, and the flexibility of cells in the oxidation of three mitochondrial fuels, namely glucose (pyruvate), glutamine (glutamate), and long-chain fatty acids, were measured using inhibitors of each metabolic pathway which were injected in a different order and combination. Figure 2G shows the three fundamental parameters for each source of energy. When we analyzed the role of glucose as an energy source, no difference was detected in terms of dependence, capacity, and flexibility between transfected cells and the control. However, when we analyzed glutamine as an energy source, inhibiting the two alternative pathways, the cells transfected with bORF3c showed a significant increase in capacity, in comparison with both cells transfected with the empty plasmid and cells overexpressing hORF3c. In addition, cells transfected with bORF3c showed an increase in flexibility, compared to cells transfected with hORF3c. These cells, therefore, seem to be able to adapt their metabolism by exploiting other fuels when the glutamine pathway is blocked by the BPTES (bis-2-(5-phenylacetamido-1,3,4-thiadiazol-2-yl) ethyl sulfide) inhibitor. On the other hand, cells overexpressing hORF3c protein displayed a slight increase in glutamine dependence compared to the control, and a significant decrease in flexibility compared to bORF3c. This result indicates that the mitochondria of these cells are unable to bypass the blocked pathway by oxidizing other fuels. When fatty acids were investigated as an energy source, cells overexpressing both ORF3c proteins exhibited a significantly higher dependence compared to the control, as shown in Figure 2G. In conclusion, the mitochondria of transfected cells were not only unable to bypass the blockage of the fatty acid pathway through the use of the other two fuels, but they also required fatty acids to maintain basal OCR.

### Hyperactivation of oxidative phosphorylation is sustained by fatty acids oxidation

Based on seahorse analysis, we investigated the role of NAD+/NADH ratio as the regulator between mitochondrial fatty acid synthesis and oxidation (Schwartz et al, 1974). In general, fatty acid β-oxidation starts in the presence of an abundant phosphate acceptor and with the consumption of NADH, which leads to an increase in the NAD+/NADH ratio. Conversely, during fatty acids synthesis the phosphate acceptor is lacking, while the substrate is present in excess, and most of the NAD+ is reduced. Measurements of NAD+/NADH ratio in transfected and control HSAEC1 cells showed that the overexpression of hORF3c protein increased NADH level, markedly decreasing NAD+/NADH ratio (Fig. 3A). A smaller, not statistically significant decrease in the ratio was also observed in cells overexpressing bORF3c (Fig. 3A). These results indicate that cells transfected with hORF3c increase not merely their use of fatty acids as carbon source, but also their rate of fatty acid synthesis to maintain the equilibrium between catabolism and anabolism. A change in NAD+/NADH ratio is only a mediator of the equilibrium between fatty acids oxidation and synthesis that are supported by the presence of Krebs Cycle substrates. In particular, succinate is the only substrate that can reduce a large pool of mitochondrial NAD+ and keep it reduced, whereas citrate could support fatty acid synthesis. Higher levels of citrate and succinate were observed after transfection with either viral proteins (Fig. 3B). At the same time, the amount of malate and alfa-ketoglutarate did not reveal any differences between samples. Only a slight increase in alfa-ketoglutarate was observed after hORF3c transfection, although not statistically significant (Fig. 3B).

**Figure 3.**
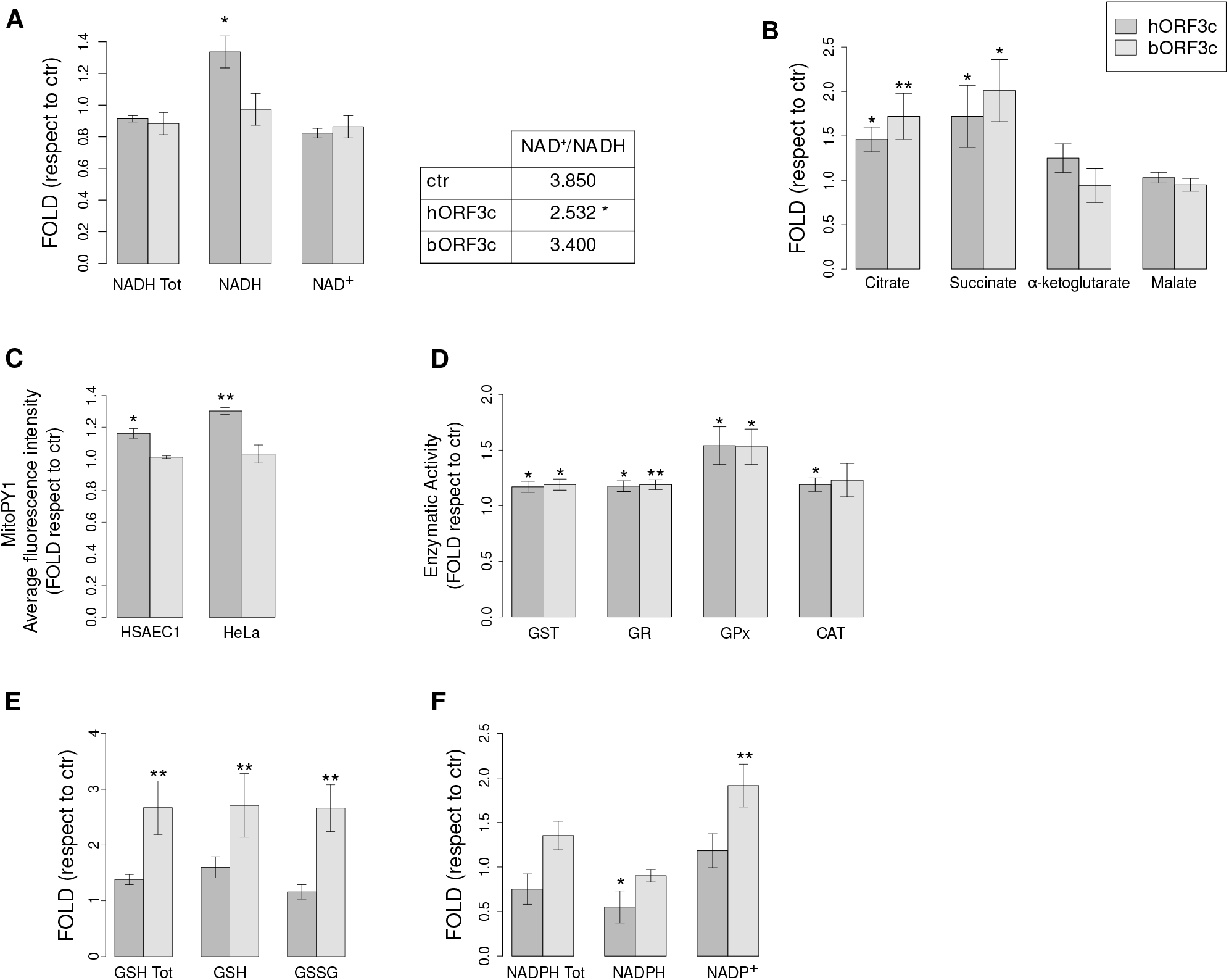
ORF3c induces oxidative stress and increases succinate levels. **(A)** NADH total, NADH and NAD+ level. In the table is reported the relative NAD+/NADH ratio. **(B)** Analysis of Krebs cycle intermediate levels in HSAEC1 cells transfected with either human or bat ORF3c. **(C)** Analysis of mitochondrial H_2_O_2_ production in HSAEC1 and HeLa cells transfected with ORF3c from either human or bat SARS-CoV-2. After incubation with 5 μM MitoPY1, the level of cell fluorescence was measured. **(D)** Activities of enzymes involved in oxidative stress defense. **(E)** Glutathione total (GSH Tot), reduced glutathione (GSH) and oxidized glutathione (GSSG) levels. **(C)** NADPH total, NADPH and NAPD+ level. Bars indicate the mean ± SEM of three biological replicates. Statistical significance: * p < 0.05, ** p < 0.01 (Dunnett’s test).

Because the increase in mitochondrial oxygen consumption due to succinate accumulation can be related to an upregulated mitochondrial subunit content, we used Real-Time PCR to investigate the level of transcripts coding for the various subunits of the five respiratory complexes. We did not detect any significant increase in the level of transcripts in transfected cells compared to the empty plasmid, with the exception of COXIII and CytB genes, showing a slight increase in expression following transfection with hORF3c (Supplementary Fig. S2C). The increase in succinate level could be linked to Reverse Electron Transport (RET) (Scialò et al, 2017; Tretter et al, 2016).

This condition allows cells to use part of the electron flow from succinate to reverse electron transfer through complex I, reducing NAD+ to NADH as shown in Figure 3A, while another part of the electron flow follows the canonical pathway from CoQ to complex IV and oxygen reduction. The hypothesis seems to be verified only in cells transfected with hORF3c because, as well as a reduction of NAD+ to NADH, saturating levels of succinate also lead to a quick conversion of ADP to ATP, and high mitochondria membrane potential, as previously shown. Moreover, the rate of ROS production, especially hydrogen peroxide (H_2_O_2_), in RET is very high (Korshunov et al, 1997).

### ORF3c expression enhances oxidative stress

To further investigate the RET hypothesis, mitochondrial hydrogen peroxide generation was measured using MitoPY1. Results showed that only the overexpression of hORF3c leads to an increase in mitochondrial H_2_O_2_ production in both HeLa and HSAEC1 cell line models (Fig. 3C).

However, in order to evaluate the effect of the overexpression of hORF3c and bORF3c proteins in the context of the oxidative stress response induced by an increase of H_2_O_2_, we assayed the activities of different antioxidant enzymes involved in ROS detoxification. In particular, except for catalase (CAT), all the enzymes shown in Figure 3D play a role in mitochondrial ROS elimination, in a direct or indirect manner. Glutathione S-transferase (GST) is involved in detoxification mechanisms via conjugation of reduced glutathione with numerous substrates; glutathione reductase (GR), a flavoprotein, catalyzes the reduction of glutathione disulfide (GSSG) to glutathione (GSH) with the participation of NADPH as an electron donor; glutathione peroxidase (GPx), a cytosolic enzyme, catalyzes the reduction of hydrogen peroxide to water and oxygen, as well as the reduction of peroxide radicals to alcohols and oxygen. Catalase, which plays its role mainly in peroxisomes, catalyzes the decomposition of hydrogen peroxide to water and oxygen. As shown in Figure 3D, the overexpression of hORF3c and bORF3c proteins leads to a significant increase in the enzymatic activity of GST, GR and GPx compared to the control; a significant increase of CAT was instead observed only in the presence of hORF3c (Fig. 3D). Although mammalian cells have evolved antioxidant enzymes to protect against oxidative stress, the most important factor in H_2_O_2_ elimination is the availability of NADPH. Indeed, this substrate is required for the regeneration of reduced glutathione, used by GPx and GST, through GR. As reported in Figure 3F a significant decrease of NADPH as reducing power was observed in the presence of hORF3c with respect to the control. Cconversely, bORF3c induced a significant increase in NADP+. Glutathione assay showed that total glutathione level was significantly higher after transfection with bORF3c (Fig. 3E).

These data support the idea that cells transfected with the hORF3c protein are not able to adequately eliminate accumulated hydrogen peroxide, whereas cells transfected with bORF3c, although showing some mild signs of oxidative stress, are able to buffer its negative effects thanks to the presence of a sufficient amount of ROS scavengers.

### SARS-CoV-2 ORF3c counteracts autophagy

Mitochondria are most commonly associated with energy production through oxidative phosphorylation, but they are also involved in a myriad of other functions, including innate immune response. Upon infection of a target cell, SARS-CoV-2 may be recognized by innate immunity sensors inducing signaling cascades that lead to the release of IFNs and pro-inflammatory cytokines, as well as to activation of autophagy for lysosomal degradation of virus/viral component (Beyer & Forero, 2022; Hayn et al, 2021). SARS-CoV-2 has evolved a wide variety of strategies to disarm innate host defenses (Beyer & Forero, 2022). For instance, it can alter mitochondrial functions leading to enhanced ROS production, perturbed signaling, and blunted host antiviral defenses. In this respect, an important role is played by accessory proteins, including ORF9b and ORF10, which, like ORF3c, have a mitochondrial localization (Han et al, 2021; Jiang et al, 2020; Li et al, 2022a).

To explore the function of ORF3c in the antiviral innate immune response, we analyzed the impact of hORF3c and bORF3c on IFN induction, IFN/pro-inflammatory cytokine and chemokine signaling, as well as on autophagy. Thus, we evaluated the release of 26 cytokines and chemokines in cells transfected with vectors expressing EGFP, hORF3c, and bORF3c. No differences were observed among plasmids. Likewise, the expression of 62 genes involved in innate and adaptive immunity was not modulated by the viral ORFs.

We next explored whether ORF3c affects autophagy, an evolutionary conserved intracellular process that delivers proteins and organelles to the lysosomes for degradation, through the formation of double-membrane vesicles, termed autophagosomes. Autophagy is also a key mechanism adopted by the host cell for clearing pathogens. To promote their survival and replication, many viruses, including SARS-CoV-2, have evolved mechanisms for interfering with the formation or maturation of autophagosomes in host cells (Koepke et al, 2021; Mao et al, 2019).

Thus, we analyzed the levels of the autophagosomal markers LC3 and p62 protein, the latter targeting poly-ubiquitinated proteins to autophagosomes for degradation, in ORF3c-transfected cells. During autophagosome formation, the cytosolic LC3-I isoform is converted into an active phosphatidylethanolamine-conjugated form, LC3-II, that is incorporated in the autophagosomal membrane. Thus, LC3-II amount is considered a reliable autophagosomal marker (Kabeya et al, 2000). Therefore, HeLa cells were transfected with vectors expressing hORF3c, bORF3c or with the control vector expressing the EGFP-DDK tag, and total protein extracts were analyzed. We found that hORF3c induced an increase in LC3-II and p62 levels (Fig. 4A) compared with the control, indicating the presence of an increased number of autophagosomes, while bORF3c did not affect the autophagosomal marker levels. Data were confirmed by immunofluorescence by using the pCMV6-MAP1LC3B-RFP vector to stain autophagosomes (Fig. 4B). Indeed, we found that, in basal conditions, cells transfected with hORF3c presented autophagosome accumulation with an increased number of RFP-LC3/p62 vesicles (Fig. 2C and 2D) compared with control and bORF3c-transfected cells. Although starvation with EBSS (Earle’s Balanced Salt Solution) induced autophagy in all transfected cells, the number of autophagosomes remained significantly higher in hORF3c-transfected cells.

**Figure 4.**
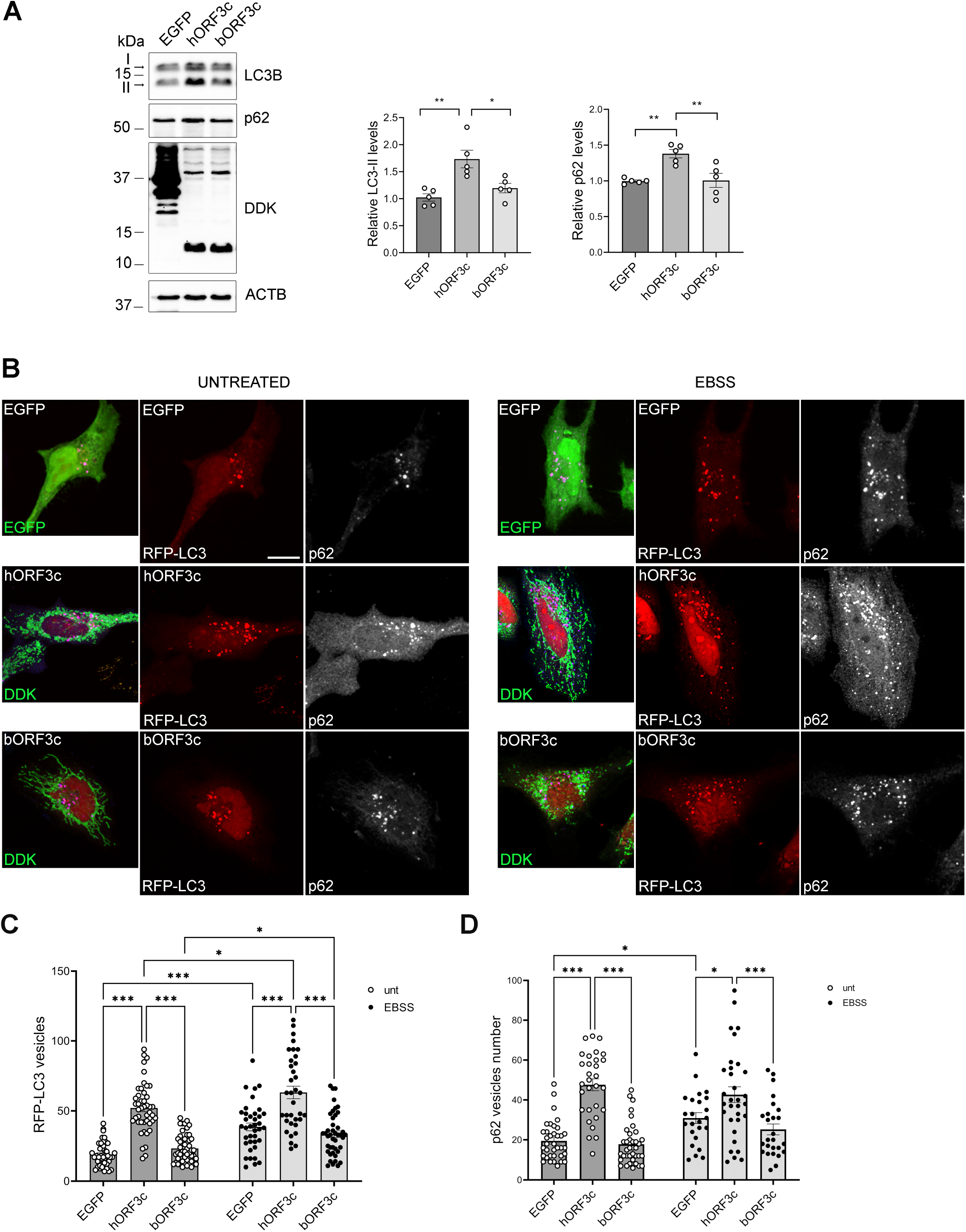
ORF3c overexpression increases autophagosome levels in cells. **(A)** HeLa cells were transfected with hORF3c, bORF3c or control vector (EGFP). Cells were lysed and total protein extracts were run onto 10/15% SDS-polyacrylamide gels and probed with anti-DDK, – LC3B, -p62/SQSTM1 and -ACTB Abs. LC3-II and p62 levels were quantified, normalized on ACTB levels and expressed as fold increase of control (one way ANOVA followed by Dunnett’s multiple comparison test; *n*=5 experiments). **(B)**. Cells were co-transfected with hORF3c, bORF3c or control vector (EGFP) and the pCMV6-MAP1LC3B-RFP vector for the staining of autophagosomes (red). After 24h, cells were starved in EBSS for 1h to induce autophagy. Treated and untreated cells were fixed and stained with anti-DDK (green) to detect ORF3c proteins, and with anti-p62 (blue) Abs. Scale bar,: 10 μm. (**C**) RFP-LC3 positive vesicles and (**D**) p62 positive vesicles are reported in the graphs (Two way ANOVA followed by Tukey’s multiple comparison test, *n*>25 cells).

An increased number of autophagosomes may derive from an increased biogenesis or from inhibition of the autophagic flux. Therefore, we analyzed autophagosome degradation by using the mRFP-GFP tandem fluorescent tagged LC3B vector to visualize autophagosomes (Fig. 5A) (Kimura et al, 2007). The GFP signal is sensitive to the acidic compartment and is quenched under low-pH conditions when autophagosomes fuse with lysosomes. We found that a very low percentage of the autophagosomes accumulated in hORF3c-transfected cells are red acidified functional autolysosomes (mRFP+, GFP-), compared with cells transfected with the control or with bORF3c (Fig. 5A). This suggests the presence of degradation defects, as reported for other SARS-CoV-2 proteins (e.g. ORF7a, ORF3a) (Hayn et al, 2021). Nevertheless, we found that the percentage of RFP-LC3 vesicles co-localizing with the lysosomal marker LAMP1 was similar in all transfected cells and in untransfected controls, suggesting that the expression of hORF3c did not affect autophagosome-lysosome fusion and that the autophagosome accumulation observed in these cells did not derive from fusion defects (Fig. 5B).

**Figure 5.**
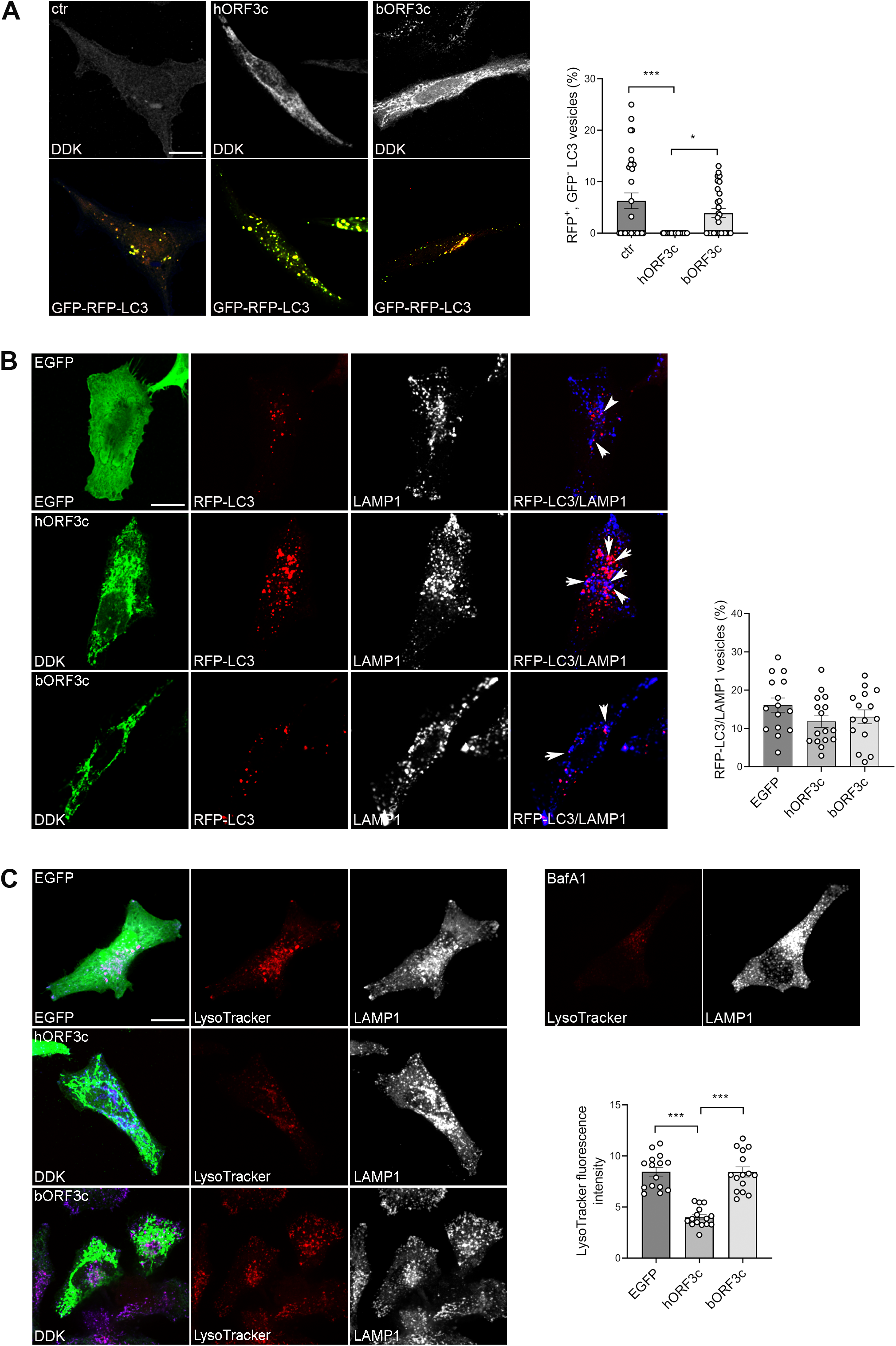
ORF3c overexpression impacts on autophagic flux. **(A)** HeLa cells were co-transfected with mRFP-GFP-LC3 and hORF3c or bORF3c or empty (ctr) vector for 24 h, fixed and stained with anti-DDK Ab. mRFP-GFP-LC3 positive autophagosomes are shown in yellow. Scale bar, 10 μm. Red mRFP^+^, GFP^-^ LC3 vesicles, corresponding to acidified autolysosomes, were counted and expressed as percentage of total LC3 vesicles (one way ANOVA followed by Dunnett’s multiple comparison test; *n*=30 cells). **(B)** Hela cells co-transfected with RFP-LC3B and hORF3c, bORF3c or EGFP vector were stained with Abs against DDK tag (green) and the lysosomal marker LAMP1 (blue). Autophagosomes (RFP-LC3) fused with LAMP1 positive vesicles were counted, normalized to total RFP-LC3 vesicles and expressed as percentage (one way ANOVA followed by Dunnett’s multiple comparison test; *n*=15 cells). **(C)** Hela cells transfected with hORF3c, bORF3c or EGFP vector were labeled with Lysotracker Red DND-99, fixed and immunostained with anti-LAMP1Ab (blue). Scale bar: 10 μm. Bafilomycin A1 (BafA1) was used as negative control. LysoTracker fluorescence intensity was quantified and reported in the graph (one way ANOVA followed by Dunnett’s multiple comparison test; *n*=15 cells).

We next assessed whether hORF3c affects lysosomal acidification by using the acidic organelle marker LysoTracker Red, a cell-permeable weak base dye which selectively accumulates in acidified vesicles, such as lysosomes and autolysosomes (Cheng et al, 2018). We observed a decrease in LysoTracker Red fluorescence intensity in hORF3c-transfected cells compared with the control, indicating a reduced acidity of lysosomes (Fig. 5C). No difference was detected between bORF3c-transfected cells and control.

In summary, these data indicate that SARS-CoV-2 ORF3c (but not bORF3c) impairs autophagy; in particular, ORF3c leads to autophagosome accumulation affecting lysosomal acidification, thus blocking the normal autophagic degradation process.

Autophagy plays an important role also in the maintenance of mitochondrial homeostasis. Indeed, the quality control of mitochondria is achieved by balanced actions among mitochondrial biogenesis, dynamics, and mitophagy, a selective autophagy that removes dysfunctional or exceeding mitochondria (Palikaras et al, 2018). Viruses often hijack mitophagy to enable immune escape and self-replication (Li et al, 2022a; Li et al, 2022b; Zhang et al, 2018). We therefore analyzed the sequestration of mitochondria in the autophagosomes in ORF3c-transfected cells by quantifying the co-localization of RFP-LC3 and the mitochondrial marker TOM20 (Supplementary Fig.S3). We did not detect differences in the percentage of mitochondria co-localizing with autophagosomes among hORF3c, bORF3c and the control (Supplementary Fig.S3). These data suggest that the ORF3c protein does not impair mitophagy.

## DISCUSSION

Coronaviruses encode a variable number of accessory proteins which differ in sequence and number even among closely related viruses. These proteins are usually dispensable for viral replication, but often play a role in host-virus interactions, in the suppression of immune responses, or in immune evasion. For these reasons, some of them represent virulence factors (Fang et al, 2021; Forni et al, 2017; Forni et al, 2022). Therefore, gaining full insight into the functions of accessory proteins is vital for understanding coronavirus pathogenesis and for developing effective antiviral drugs and vaccines. Since the beginning of the pandemic, the accessory proteins encoded by SARS-CoV-2 have been an object of study and their role in the processes of escape from the innate immune system, as well as their interaction with host proteins, have been reported. Although highly conserved in sarbecoviruses and considered a protein with potential functional role (Cagliani et al, 2020; Finkel et al, 2021; Firth, 2020; Jungreis et al, 2021), the accessory protein ORF3c of SARS-CoV-2, an alternative open reading frame within the *ORF3a* gene, attracted little attention. To cover this gap, we characterized ORF3c in terms of cellular localization, host’s antiviral response modulation, and effects on mitochondrial metabolism. Our data show that ORF3c has a mitochondrial localization, alters mitochondrial metabolism, and increases ROS production. Conversely, we did not observe an action of the protein on the modulation of the expression of interferons and cytokines/chemokines. ORF3c also acts on autophagy blocking the autophagic flux and inducing the accumulation of autophagosomes/autolysosomes. In particular, our data suggested that ORF3c expression prevents autophagic degradation by altering the pH of lysosomes.

Several RNA viruses can induce or manipulate the autophagic responses, exploiting autophagosomes to facilitate viral replication and to elude the innate immune response (Wong & Sanyal, 2020). Among these, SARS-CoV-2 restricts autophagy-associated signaling and blocks autophagic flux. In particular, cells infected with SARS-CoV-2 show an accumulation of key metabolites, the activation of autophagy inhibitors, and a reduction in the levels of several proteins responsible for processes spanning from autophagosome formation to autophagosome–lysosome fusion and lysosome deacidification (Gassen et al, 2021; Ghosh et al, 2020).

Recently, different studies analyzed the effect of individual SARS-CoV-2 proteins on autophagy and identified several viral proteins involved in this process. Some of them act by causing an increase or inhibition in autophagy, but most of the viral proteins (E, M, ORF3a and ORF7a) promote the accumulation of autophagosomes, also reducing autophagic flow (Hayn et al, 2021; Li et al, 2021). Specifically, ORF3a and ORF7a were reported to block autophagy by interfering with autophagosome-lysosome fusion and lysosomal acidification (Chen et al, 2021a; Hayn et al, 2021; Hou et al, 2022; Miao et al, 2021).

Considering all these data, we suggest that, during SARS-CoV-2 infection, various autophagy regulation mechanisms are put in place, in order to achieve a state of equilibrium that both allows inhibition of the innate immune response and favors viral replication. Different viral proteins might differently affect autophagy, resulting in a fine regulation of this pathway.

In this context, an important role is likely to be played by ORF3c, not only in SARS-CoV-2, but probably also in all sarbecoviruses, where ORF3c is highly conserved. To test this hypothesis, we evaluated the effect on autophagy of the ORF3c protein encoded by one of the bat betacoronaviruses most closely related to SARS-CoV-2 (bat-CoV RaTG13, bORF3c). In most analyses, a similar trend as that observed for SARS-CoV-2 ORF3c was evident for bORF3c, but the effect was definitely weaker. The two viral proteins (hORF3c and bORF3c) differ only in two amino acids, so we must conclude that the two substitutions in position 36 and 40 are sufficient to determine a different effect on autophagy. We cannot however exclude that the different action observed for bORF3c is at least partially explained by the use of human cell lines.

An important role in autophagy is played by reactive oxygen species (ROS). An increase in ROS has been described in several physiological and pathological conditions including aging, cancer, diabetes, neurodegenerative disorders, and infection (Foo et al, 2022). In most of these cases, high levels of mitochondrial ROS compromise lysosomal acidity and autophagic flow. Recently, it has been demonstrated that an increase in ROS levels in glucose-deprived fibroblasts can reduce lysosome acidification and impair autolysosome degradation, blocking the autophagic flux (Song & Hwang, 2020). Indeed, increased ROS levels could inactivate the vacuolar ATPase (vATPase), a proton pump that generates an acidic pH in the lysosome (Song & Hwang, 2020). Notably, in pulmonary cell lines overexpressing ORF3c, we found a decrease in the level of basal glycolysis and significant increase of mitochondrial hydrogen peroxide (H_2_O_2_, a non-radical ROS), oxidative phosphorylation (OXPHOS) and β-oxidation of fatty acids. The activation of these last two processes is known to induce oxidative stress (Quijano et al, 2016; Scialò et al, 2017; Zhao et al, 2019). Therefore the alteration of mitochondrial metabolism we observed in ORF3c-transfected cells could be responsible of autophagosomes/autolysosomes accumulation and lysosomes deacidification, as already reported in glucose-deprived fibroblasts (Song & Hwang, 2020). Thus, we suggest that ORF3c acts by mimicking a condition of glucose starvation. The metabolic rearrangement induced by ORF3c parallels what happens in the second phase of SARS-CoV-2 infection. Initially, in the first phase of infection, the energy supply occurs mainly through hyperactivation of glycolysis, which culminates with the reduction of pyruvate into lactate. On the other hand, mitochondrial oxidative phosphorylation is very marginal to energy production: the respiratory complexes allow an electron transfer with poor efficiency, and the electrochemical potential across the inner mitochondrial membrane is low. This first phase is functional for the replication of the virus and its expansion in the host organism.

The second phase is a chronic degeneration of cellular physiology (Shenoy, 2020); at this point, in line with what happens by transfecting cells with the accessory protein ORF3c, oxidative phosphorylation is the main way of energy production, glycolysis being downregulated. Fatty acids become the primary energy substrate, beta-oxidation being upregulated; glucose consumption and lactate production decrease, reducing acidification. Acetyl-CoA is channeled into the citrate cycle, which proceeds predominantly in the canonical direction. There is a shift from glucose oxidation to fatty acids oxidation.

How ORF3c achieves this result remains unclear and further studies are required to establish in detail the mechanism by which the viral protein alters mitochondrial metabolism and blocks the autophagic flux. Likewise, it is unclear how ORF3c can alter the metabolic state of infected cells. Given its mitochondrial localization, we hypothesize that the ORF3c protein does not act directly on the glycolytic process, but rather on the transport of pyruvate from the cytoplasm to the mitochondrial matrix or in the early stages of pyruvate modification. Specific experiments to answer these questions will be necessary to fully understand the role played by the ORF3c protein during SARS-CoV-2 infection.

## MATERIALS AND METHODS

### Protein structure prediction

The three-dimensional structures of SARS-CoV-2 and RaTG13 ORF3c proteins were predicted using the Robetta online protein structure prediction server (https://robetta.bakerlab.org/) (Baek et al, 2021). Robetta can predict the three-dimensional protein structure given an amino acid sequence. The default parameters were used to produce models using the simultaneous processing of sequence, distance, and coordinate information by the three-track architecture implemented in the RoseTTAfold method (Baek et al, 2021). For both proteins, the confidence of the model was good (*Global Distance Test, GTD*, > 0.5). 3D structures were rendered using PyMOL (The PyMOL Molecular Graphics System, Version 1.8.4.0; Schrödinger, LLC). The predicted structural model 1 of the top five models of both proteins were used to perform the structural superposition, using the align command. The RMSD value was also calculated with PyMOL.

### Plasmids

Complementary DNA (cDNA) containing coding sequences of ORF3c sequence of SARS-CoV-2 (hORF3c, NC_045512.2, nucleotide position: 25457-25579) and RatG13 (bORF3c, MN996532, nucleotide position: 25442-25564) were synthesized by Origene custom service. The hORF3c and bORF3c were cloned in pCMV6-Entry Mammalian Expression Vector (Origene, PS100001) in frame with C-terminus Myc-DDK tag. Likewise, EGFP was cloned in pCMV6-Entry (pCMV6-EGFP). The commercial expression vectors pDsRed2-Mito (Clontech Laboratories, Inc., CA, USA), pCMV6-RFP-MAP1LC3B (Origene, RC100053) were used for fluorescent labeling of mitochondria and autophagosomes, respectively. To analyse autophagosome degradation, cells were transfected with the mRFP-GFP-LC3 (ptfLC3) vector, a gift from Tamotsu Yoshimori (Addgene plasmid #21074) (Kimura et al, 2007).

### Cell lines and culture conditions

Human epithelial adenocarcinoma HeLa (ATCC, CCL-2) cells and human epithelial lung carcinoma A549 (ATCC, CCL-185) cells were cultured in Dulbecco’s Modified Eagle’s Medium (DMEM, Euroclone, Milano, Italy) supplemented with 10% Fetal Bovine Serum (FBS, Euroclone, Milano, Italy), 2 mM L-glutamine and 100 U/ml penicillin/streptomycin (Invitrogen, Carlsbad, CA, USA, Thermo Fisher Scientific, Waltham, MA, USA). The normal human lung cell line HSAEC1-KT (ATCC^®^ CRL-4050™) was grown in SABM Basal Medium™ supplemented with Bovine Pituitary Extract (BPE), Hydrocortisone, human Epidermal Growth Factor (hEGF), Epinephrine, Transferrin, Insulin, Retinoic Acid, Triiodothyronine, Bovine Serum Albumin – Fatty Acid Free (BSA-FAF), 100 U/ml penicillin and 100 μg/ml streptomycin. All the reagents for HSAEC1 cell culture were supplied by Lonza (Lonza Group, Basel, Switzerland). Cell lines were maintained at 37°C in a humidified 5% CO_2_ incubator. All cell lines were tested for mycoplasma contamination (MP0035; Merck Life Science).

Autophagy was induced by amino acid and serum starvation in Earle’s Balanced Salt Solution (EBSS, ECB4055L, Euroclone) for the indicated times.

### Immunostaining and confocal immunofluorescence

HeLa/A549/HSAEC1 cells were seeded (0.3 x 105 cells/well) 24 h before transfection into 6-well plates onto coverslips treated with 0.1 ug/mL poly-L-lysine. Transient transfections were performed using Lipofectamine 2000 (Thermo Fisher Scientific, Waltham, MA, USA) with 2.5 μg of plasmid DNA (pCMV6-hORF3c, pCMV6-bORF3c, pCMV6-Entry, pCMV6-EGFP), according to manufacturer’s instruction. For the staining of autophagosomes and mitochondria, cells were cotransfected with the pCMV6-RFP-MAP1LC3B vector and with the pDsRed2-Mito vector, respectively. Co-trasfections were performed with 2 μg of each plasmid. At 24 hours after transfection, cells were fixed with 4% paraformaldehyde (Santa Cruz Biotechnology, sc-281692) and permeabilized with phosphate-buffered saline (PBS; Euroclone, ECB4053L) containing 0.1% saponin (Merck Life Science, S4521) and 1% BSA (Merck Life Science, A9647). Samples were then incubated for 2 h with primary antibodies and revealed using the secondary antibodies Alexa Fluor 488, 546 and 647 (Invitrogen, Thermo Fisher Scientific). Nuclei were stained with DAPI. To analyse autophagosome degradation, cells were transfected with the mRFP-GFP-LC3 (ptfLC3) vector, fixed with cold methanol for 5 min and permeabilized with PBS containing 0.1% Triton X-100 (Merck Life Science, T8787). For the staining of acidic organelles, cells were incubated with 75 nM LysoTracker Red DND-99 (L7528, Invitrogen, Thermo Fisher Scientific) for 5 minutes to avoid alkalinization, accordingly with manufacturer instructions, fixed in paraformaldehyde and processed. The list of antibodies is reported in Supplementary Table 1.

Confocal microscopy was performed with a Yokogawa CSU-X1 spinning disk confocal on a Nikon TiE inverted microscope equipped with a Nikon 60x/1.40 oil Plan Apochromat objective and were acquired with an Andor Technology iXon3 DU-897-BV EMCCD camera (Nikon Instruments S.p.A., Firenze, Italy). RFP-LC3, p62 and LAMP1 positive vesicles were counted with ImageJ/Fiji by using the “analyze particles” tool and the investigator was blinded as to the nature of the sample analyzed.

### Mitochondria isolation and fractionation

HeLa cells were seeded (1.2 x 106 cells/well) into p100 plates 24 h before transfection. Transient transfections were performed using Lipofectamine™ 3000 Transfection Reagent (Thermo Fisher Scientific, Waltham, MA, USA) with 15 μg of plasmid DNA/plate (pCMV6-hORF3c and pCMV6-bORF3c), according to the manufacturer’s instruction. 24 h post transfection cells were rinsed twice with PBS and harvested by centrifugation. Mitochondria isolation was performed using the Mitochondria Isolation Kit for Cultured Cells (Thermo Fisher Scientific, Waltham, MA, USA) using the reagent-based method starting from about 2 x 10^7^ cells for each construct, according to the manufacturer’s protocol. For each sample, total extracts were fractionated, separating intact mitochondria from cytosol. After isolation, mitochondria were lysed with 2% CHAPS in 25mM Tris, 0.15M NaCl, pH 7.2 and centrifuged at high speed to separate the soluble fraction (supernatant) to the insoluble fraction (pellet).

### Co-immunoprecipitation assays

Co-immunoprecipitation assays were performed with the Pierce™ MS-Compatible Magnetic IP Kit, protein A/G (Thermo Fisher Scientific, Waltham, MA, USA). Briefly, 24 h post transfection HeLa cells were rinsed twice with ice-cold PBS and lysed on ice in IP-MS Cell Lysis Buffer added of Halt™ Protease Inhibitor Cocktail EDTA-free (Thermo Fisher Scientific, Waltham, MA, USA), for 10 minutes with periodic mixing. Extracts were clarified by centrifugation (13,000 × g for 10 minutes) and quantified by Pierce™ BCA Protein Assay Kit (Thermo Fisher Scientific, Waltham, MA, USA). 500 μg of cell lysate were combined with 5μg of IP antibody and incubated overnight at 4°C with mixing to form the immune complex. The immunoprecipitation reaction was performed for 1h at RT, by incubating the sample/antibody mixture with 0.25mg of pre-washed Pierce Protein A/G Magnetic Beads. After washes, target antigen samples were eluted in IP-MS Elution Buffer and dried in a speed vacuum concentrator. Samples were reconstituted in Sample Buffer for SDS-PAGE/WB analyses.

### SDS-PAGE and Western blotting

After 24h post transfection, cells were rinsed with ice-cold PBS, harvested by scraping and lysed in Lysis buffer (125 mM Tris/HCl pH 6.8, 2.5% SDS). Lysates were incubated for 2 min at 95°C. Homogenates were obtained by passing 5 times through a blunt 20-gauge needle fitted to a syringe and then centrifuged at 12,000xg for 8 min. Supernatants were analyzed for protein content by Pierce™ BCA Protein Assay Kit (Thermo Fisher Scientific, Waltham, MA, USA). SDS-PAGE and Westernblot were carried out by standard procedures: samples were loaded and separated on a 10%, 12% or 15% acrylamide/bis-acrylamide SDS-PAGE, blotted onto a nitrocellulose membrane (Amersham, Cytiva, Marlborough, MA, USA). Horseradish peroxidase-conjugated secondary antibodies were used and signals were detected using ECL (GE Healthcare) and acquired with iBrightFL1000 (Thermo Fisher Scientific) (Supplementary Table 1). Protein levels were quantified by densitometry of immunoblots using Fiji ImageJ software.

### Quantigene Plex gene expression assay

Gene expression analysis was performed on 50.000 A549-ACE2 transfected cells by QuantiGene Plex assay (Thermo Scientific, Waltham, MA, USA). This approach provides a fast and high-throughput solution for multiplex gene expression quantitation, allowing for the simultaneous measurement of 64 custom-selected genes in a single well of a 96-well plate. Host genes were selected based on their involvement in human coronavirus infections. The QuantiGene Plex assay is hybridization-based and incorporates branched DNA technology, which uses signal amplification for direct measurement of RNA transcripts. The assay does not require RNA purification, nor retro-transcription, with a minimal sample handling. Some of the targets resulted below the detection limit and the arbitrary value of 0 was assigned. Results were calculated relative to *GAPDH, β-Actin* and *PPIB* as housekeeping genes, and expressed as ΔCt. The list of host genes is reported in Supplementary Table 2.

### Cytokine and Chemokine Measurement by Multiplex Assay

Cytokine/chemokine concentration was assessed in cell culture supernatants 48 h post transfection by using immunoassays formatted on magnetic beads (Bio-Plex Pro Human Cytokine 27-plex Assay #M500KCAF0Y) (Bio-Rad, Hercules, CA, USA), according to the manufacturer’s protocol via Luminex 100 technology (Luminex, Austin, TX, USA). For the targets over-range an arbitrary value of 4000 pg/mL is assigned, while 0 pg/mL is attributed to values below the detection limit. The list of cytokines/chemokines is reported in Supplementary Table 2.

### Viability assay

In order to evaluate the effect of ORF3c from SARS-CoV-2 or from BatCov RaTG13 on cell viability, HSAEC1 cells were seeded in 96-well plates at a density of 1 × 10^4^ cells/well and after 24 h were transiently transfected using Lipofectamine 2000 (Thermo Fisher Scientific, Waltham, MA, USA). After an incubation at 37°C for 36 h post transient transfection, the medium was replaced with complete medium without phenol red and 10 μL of 5 mg/mL MTT solution (In vitro toxicology assay kit, MTT-based, TOX-1KT, Merck, Darmstadt, Germany) were added to each well. After a further 4 h incubation time, absorbance upon solubilization was measured at 570 nm using a micro plate reader. Viabilities were expressed as a percentage of the mock (pCMV6-vector). No effect on cell viability was detected.

### Oxygen consumption rate and extra-cellular acidification rate measurements

Oxygen consumption rate (OCR) and extra-cellular acidification rate (ECAR) were investigated using Agilent Seahorse XFe96 Analyzer on HSAEC1 cell line transfected with ORF3c from SARS-CoV-2 or ORF3c from BatCov RaTG13.

Cells were seeded in Agilent Seahorse 96-well XF cell culture microplates at a density of 4 × 104 cells per well in 180 μL of growth medium and after 24 h were transiently transfected.

Before running the assay, the Seahorse XF Sensor Cartridge was hydrated and calibrated with 200 μL of Seahorse XF Calibrant Solution in a non-CO2 37 °C incubator to remove CO2 from the media that would otherwise interfere with pH-sensitive measurements.

After an incubation at 37°C for 36 h post transient transfection, the growth medium was replaced with 180 μL/well of Seahorse XF RPMI Medium, pH 7.4 with 1 mM Hepes, without phenol red, containing 1 mM pyruvate, 2 mM L-glutamine and 10 mM glucose. Subsequently, the plate was incubated into a 37 °C non-CO2 incubator for 1 hour, before starting the experimental procedure, and the compounds were loaded into injector ports of the sensor cartridge.

For Agilent Seahorse XF Cell Mito Stress Test Kit, pre-warmed oligomycin, FCCP, rotenone and antimycin A were loaded into injector ports A, B and C of sensor cartridge at a final working concentration of 1 μM, 2 μM and 0.5 μM, respectively. OCR and ECAR were detected under basal conditions followed by the sequential addition of the compounds and non-mitochondrial respiration, maximal respiration, proton leak, ATP respiration, respiratory capacity and coupling efficiency can be evaluated.

For Agilent Seahorse XF Glycolytic Rate Assay Kit, pre-warmed combination of rotenone and antimycin A at working concentration of 0.5 μM and 2-deoxy-D-glucose (2-DG) at 50 mM were loaded into injector ports A and B, respectively. OCR and ECAR were detected under basal conditions followed by the sequential addition of the compounds to measure basal glycolysis, basal proton efflux rate, compensatory glycolysis and post 2-DG acidification.

Using the Agilent Seahorse XF Mito Fuel Flex Test Kit, the mitochondrial fuel consumption in living cells was determined and, through OCR measuring, the dependency, capacity and flexibility of cells to oxidize glucose, glutamine and long-chain fatty acids can be calculated. Pre-warmed working concentration of 3 μM BPTES, 2 μM UK5099 or 4 μM etomoxir were loaded into injector port A and compounds mixture of 2 μM UK5099 and 4 μM etomoxir, 3 μM BPTES and 4 μM etomoxir or 3 μM BPTES and 2 μM UK5099 into injector port B to determine glutamine, glucose and long-chain fatty acid dependency, respectively. On the contrary, fuel capacity is measured by the addition into injector port A of 2 μM UK5099 and 4 μM etomoxir, 3 μM BPTES and 4 μM etomoxir or 3 μM BPTES and 2 μM UK5099 working concentration, followed by injection in port B of 3 μM BPTES, 2 μM UK5099 or 4 μM etomoxir working concentration for glutamine, glucose and long-chain fatty acid, respectively. All kits and reagents were purchased from Agilent Technologies (Santa Clara, CA, USA).

### Enzymatic activities and metabolite assays

After 36 h post transfection, HSAEC1 cells were rinsed with ice-cold PBS, harvested by scraping and lysed in 50 mM Tris-HCl, pH 7.4, 150 mM NaCl, 5 mM EDTA, 10 % glycerol, 1 % NP40 buffer, containing 1 μM leupeptin, 2 μg/mL aprotinin, 1 μg/mL pepstatin and 1 mM phenylmethylsulfonyl fluoride (PMSF). After lysis on ice, homogenates were obtained by passing the cells 5 times through a blunt 20-gauge needle fitted to a syringe and then centrifuging at 15,000g for 30 min at 4°C. Enzyme activities were assayed on supernatants. Lactate dehydrogenase (LDH) was evaluated measuring the disappearance of NADH at 340 nm according to Bergmeyer (Bergmeyer, 1974). The protein samples were incubated with 85 mM potassium phosphate buffer, 0.2 mM NADH, 0.6 mM pyruvate. Glutathione S-transferase (GST) was measured as reported in Habig (Habig et al, 1974), using 1 mM reduced glutathione (GSH) and 1 mM 1-chloro-2,4-dinitrobenzene (CDNB) as substrates in the presence of 90 mM potassium phosphate buffer pH 6.5. The reaction was monitored at 340 nm. Glutathione reductase (GR) was measured following the disappearance of NADPH at 340 nm according to Wang (Wang et al, 2001). The protein samples were incubated with 100 mM potassium phosphate buffer pH 7.6, 0.16 mM NADPH, 1 mM EDTA, 1 mg/mL BSA, 4.6 mM oxidized glutathione (GSSG). The glutathione peroxidase (GPx) activity is based on the oxidation of GSH using H_2_O_2_ as substrate, coupled to the disappearance of NADPH by glutathione reductase (GR), according to Nakamura (Nakamura et al, 1974). The protein samples were incubated with 50 mM sodium phosphate buffer pH 7.5, 0.16 mM NADPH, 1 mM NaN3, 0.4 mM EDTA, 1 mM GSH, 0.2 mM H_2_O_2_, 2 U/mL GR. Catalase (CAT) activity was evaluated according to Bergmeyer (Bergmeyer, 1983), using 12 mM H_2_O_2_ as substrate in the presence of 50 mM sodium phosphate buffer, pH 7.5. The reaction was monitored at 240 nm.

L-citrate, L-succinate, α-ketoglutarate, L-malate, NAD+/NADH, NADP+/NADPH were evaluated using kits based on colorimetric assays (Citrate Assay Kit, MAK057; Succinate Colorimetric Assay Kit, MAK184; α-ketoglutarate Assay Kit, MAK054; Malate Assay Kit, MAK067; NAD/NADH Quantitation kit, MAK037; NADP/NADPH Quantitation kit, MAK038; Merck, Darmstadt, Germany). For glutathione detection, cells were trypsinized and harvested by centrifugation at room temperature, for 10 min at 1,200×g. Pellets were washed in 3 mL PBS, harvested by a centrifugation and weighed to normalize the results to mg of cells. Pellets were resuspended in 500 μL cold 5% 5-sulfosalicylic acid (SSA), lysed by vortexing and by passing through a blunt 20-gauge needle fitted to a syringe 5 times. All the samples were incubated for 10 min at 4 °C and then centrifuged at 14,000×g for 10 min at 4 °C. The supernatant was prepared and used for the analysis following the instructions of Glutathione Colorimetric Detection Kit (catalog number EIAGSHC, Invitrogen, Carlsbad, CA, USA). The Kit is designed to measure oxidized glutathione (GSSG), total glutathione (GSH tot) and reduced glutathione (GSH tot—GSSG) concentrations through enzymatic recycling assay based on glutathione reductase and reduction of Ellman reagent (5,5-dithiobis(2-nitrobenzoic acid)) and using 2-vinylpyridine as reagent for the derivatization of glutathione (Griffith, 1980). Therefore, it was possible to obtain GSH/GSSG ratio, a critical indicator of cell health. The absorbance was measured at 405 nm using a micro plate reader. The values of absorbance were compared to standard curves (GSH tot and GSSG, respectively) and normalized to mg of cells. Final concentrations were expressed in nmol/mg cells.

### Detection of mitochondrial hydrogen peroxide

MitoPY1 (Tocris Bioscience, Bristol, UK) indicator was used to detect the mitochondrial hydrogen peroxide production in intact adherent cells. The oxidation of this probe forms intermediate probe-derived radicals that are successively oxidized to generate the corresponding fluorescent products(Winterbourn, 2014). HSAEC1 and HeLa cells were seeded in 96-well plates at a density of 1 × 104 cells/well and after 24 h were transiently transfected. After an incubation at 37°C for 36 h post transient transfection, the cells were stained with MitoPY1 at 5 μM final concentration in 1 PBS for 20 min in the dark at 37 °C. After staining, the cells were washed by warm PBS and the fluorescence (λem = 485 nm/λex = 528 nm) was measured using a fluorescence microtiter plate reader (VICTOR X3) and analyzed by the PerkinElmer 2030 Manager software for Windows.

### Mitochondrial transmembrane potential (MTP) assay

MTP alterations were assayed through fluorescence analysis, using the green fluorescent membrane dye 3,3’-dihexyloxacarbocyanine Iodide (DiOC6), which accumulates in mitochondria due to their negative membrane potential and can be applied to monitor the mitochondrial membrane potential. After 36 h post transfection, cells were incubated with 40 nM DiOC6 diluted in PBS for 20 min at 37 °C in the dark and rinsed with PBS; after adding PBS, fluorescence was measured (excitation = 484 nm; emission = 501 nm) using VICTOR Multilabel plate reader (PerkinElmer, Waltham, MA, USA).

### RNA isolation and Q-PCR

Total RNA was isolated from cells using RNeasy Mini Kits (Qiagen, Chatsworth, CA, USA), according to the manufacturer’s instructions. RNA was reverse-transcribed using SuperScript^®^ II RT (Invitrogen, Carlsbad, CA, USA), oligo dT and random primers, according to the manufacturer’s protocol.

For quantitative real-time PCR (Q-PCR), SYBR Green method was used. Briefly, 50 ng cDNA was amplified with SYBR Green PCR Master Mix (Applied Biosystems, Foster City, CA, USA) and specific primers (100 nM), using an initial denaturation step at 95°C for 10 min, followed by 40 cycles of 95°C for 15 sec and 59°C annealing for 1 min. Each sample was analyzed for NADH dehydrogenase subunit 2 (ND2), cythchrome b (cyt b), cytochrome c oxidase subunit I (COX I), cytochrome c oxidase subunit II (COX II), cytochrome c oxidase subunit III (COX III), ATP synthase F0 subunit 6 (ATP6) and ATP synthase F0 subunit 8 (ATP8) expression and normalized for total RNA content using β-actin gene as an internal reference control. The relative expression level was calculated with the Livak method (2[-ΔΔCt]) and was expressed as a fold change ± standard deviation. The accuracy was monitored by the analysis of melting curves. The following primers were used: ND2 Fw 5’-CCAGCACCACAACCCTACTA-3’ and Rv 5’-GGCTATGATGGTGGGGATGA-3’; cyt b Fw 5’-TGAAACTTCGGCTCACTCCT-3’ and Rv 5’-CCGATGTGTAGGAAGAGGCA-3’; COX I Fw 5’-GAGCCTCCGTAGACCTAACC-3’ and Rv 5’-TGAGGTTGCGGTCTGTTAGT-3’; COX II Fw 5’-ACCGTCTGAACTATCCTGCC-3’ and Rv 5’-AGATTAGTCCGCCGTAGTCG-3’; COX III Fw 5’-ACCCACCAATCACATGCCTA-3’ and Rv 5’-GTGTTACATCGCGCCATCAT-3’; ATP6 Fw 5’-GCCACCTACTCATGCACCTA-3’ and Rv 5’-CGTGCAGGTAGAGGCTTACT-3’; ATP8 Fw 5’-TGCCCCAACTAAATACTACCGT-3’.

### Statistics

Double-blind experiments were performed. We report no data exclusion. One way ANOVA or two way ANOVA followed by Dunnett’s, Tukey’s or Sidak’s multiple comparisons tests were performed using GraphPad Prism version 9.3.0 for Windows, GraphPad Software, San Diego, California USA. Results are reported as individual data plus the mean ± SEM; *n* represents individual data, as indicated in each figure legend. *p* values of less than 0.05 were considered significant. Individual *p* values are indicated in the graphs (\**p*<0.05; \*\**p*<0.01; \*\*\**p*<0.001). Statistics is reported in each figure legend.

## Data availability

This study includes no data deposited in external repositories.

## Acknowledgements

This work was supported by the Italian Ministry of Health (“Ricerca Corrente 2022” to MS), by Fondazione Cariplo (grant CORONA, n. 2020-1353), and by Regione Lombardia (Bando Progetti Ricerca Covid 19 – H44I20000470002).

## Author contributions

Alessandra Mozzi: Formal analysis, Investigation, Visualization, Writing – review & editing; Monica Oldani: Formal analysis, Investigation, Visualization, Writing – review & editing; Matilde E. Forcella: Formal analysis, Investigation, Visualization, Project administration, Writing – review & editing; Chiara Vantaggiato: Formal analysis, Investigation, Visualization, Writing – review & editing; Gioia Cappelletti: Investigation; Chiara Pontremoli: Investigation; Francesca Valenti: Investigation; Diego Forni: Formal analysis, Funding acquisition, Writing – review & editing; Mara Biasin: Formal analysis, Writing – review & editing; Manuela Sironi: Conceptualization, Funding acquisition, Supervision, Writing – review & editing; Paola Fusi: Conceptualization, Supervision, Writing – original draft, Writing – review & editing; Rachele Cagliani: Conceptualization, Formal analysis, Project administration, Supervision, Writing – original draft, Writing – review & editing.

## Conflict of interest

The authors declare that they have no conflict of interest.

